# Counterbalance of Foxp3 and IDO expression at different tumor stages in aggressive breast cancer subtypes

**DOI:** 10.1101/2021.08.23.457395

**Authors:** Romina Canzoneri, Ezequiel Lacunza, Martín E. Rabassa, Fiorella A. Cavalli, Valeria Ferretti, Luis A. Barbera, Aldo Cretón, Maria Virginia Croce, Marina T. Isla Larrain

## Abstract

Foxp3 and IDO1 are known immunomodulatory molecules involved in tumor escape and could be related to tumor infiltrating lymphocytes (TILs) in the tumor microenvironment. In this study, tumoral Foxp3 and IDO1 expression in breast cancer were evaluated in relation to lymphocyte biomarkers such as CD8 and CD45R0, regulatory T cells, as well as intratumoral and stromal TILs (iTILs and sTILs, respectively). Clinical and histopathological features were also included in the analysis. Foxp3 and IDO1 were found in tumor cells showing mainly cytoplasmic patterns in 60% and 62% tumor samples, respectively. TILs were found in 76% of samples; iTILs were detected in 92% of those samples and sTILs in 55%. Foxp3+ TILs were detected only in 12% of TILs+ samples associated with tumoral Foxp3 expression. Tumoral Foxp3 was mainly expressed at lower tumor stages while IDO1 expression was associated with advanced tumor stages; both correlated with CD8+ TILs which were observed in 77% of TILs+ samples. CD45R0+ were observed in 81% of TILs+ samples and correlated with higher tumor stages and poorly differentiated tumors. In ER negative tumors, an inverse correlation between Foxp3 and IDO1 tumoral expression was found in relation to tumor stage. TNBC subtype showed a positive correlation with the presence of iTILs. *In silico* analysis showed that Foxp3 and coexpressed genes in breast cancer were associated with immune response genes. Foxp3 was found predominantly in Basal and Her2-enriched subtypes in relation to Luminal A subtype, by RNA seq and RNA microarray database analysis. In conclusion, the expression of Foxp3 and IDO1 in tumors at different stages suggests a potential compensatory mechanism to evade the strong CTL response observed. This is relevant since the cumulative data indicates that Foxp3 as well as IDO1 could be potential targets of immunotherapy in patients with tumors at different stages and for the most aggressive breast cancer subtypes such as TNBC and Her2-enriched.

**HIGHLIGHTS:**

- Foxp3 and IDO1 were found overexpressed in the cytoplasm of tumor cells, observed in BC samples at early or advanced tumor stages, respectively, both associated with abundant cytotoxic CD8+ lymphocytes infiltrates.
- Tumor immunosuppression by Foxp3 and IDO1 tumoral expression could counteract the attempt of the host’s immune system to eradicate the tumor by CD8+ and CD45R0+ TILs cytotoxic activity.
- It is well-known the tolerogenic role of Foxp3 in Treg and less understood the role of Foxp3 in tumors. However, there is increasing evidence that Foxp3+ tumor cells appear to promote an immunosuppressive microenvironment similar to Treg. Whether this is a consequence of the effect of other immunosuppressive factors secreted by the tumor cells which can be induced by Foxp3 expression, such as TGFβ, remains to be determined.

## INTRODUCTION

Breast cancer (BC) development and dissemination are influenced by intrinsic molecular heterogeneity as well as the relation between effector and suppressive immune responses [1]. The study of these mechanisms is critical for the development of new therapeutic strategies. There are three phases in the interaction between the immune system and tumors: elimination, equilibrium and escape [2]. During the elimination phase, the host immune system can destroy the transformed cells although some tumor cells may survive due to the loss of antigens or defects in antigen presentation; triggering tumor edition. In the equilibrium phase, there is a balance between anti-tumor cytokines (IL-12, IFN-γ) and those that promote tumor growth (IL-10, IL-23). The third phase of this process, also called evasion, is characterized by proliferation of tumor cells and generation of an immunosuppressant microenvironment favoring tumor growth, spread, and metastasis through the production of IL-10, TGFβ, Vascular Endothelial Growth Factor (VEGF), Indoleamine-2,3-dioxygenase (IDO) and Programmed Death-ligand 1 (PD-L1) [3].

The expression of immunomodulatory molecules in the tumor microenvironment is critical in the dissemination of malignant cells and, in consequence, in the prognosis of patients with cancer. In this sense, IDO plays a key role, catalyzing the limiting step of the tryptophan (Trp) degradation through the kynurenine (Kyn) pathway. In tumors, the expression of this enzyme correlates with advanced stages and nuclear grades as it was previously reported [4, 5]. IDO was originally described in the placenta and associated with materno-fetal tolerance; it has been implicated in protection mechanisms to graft rejection and treatment of autoimmune disorders [6-8]. The immunosuppressive effect of IDO is mediated by the host’s immunesurveillance restriction by the inhibition of activated T cells, enhancing the activity and the induction of CD4+ CD25+ Foxp3+ regulatory T cells (Treg) [9-11]. IDO inhibitors cooperate with cytotoxic agents to allow regression of tumors refractory to single therapy agents in a BC model suggesting an improvement in the chemotherapy response [12]. The presence of tumor infiltrating lymphocytes (TILs) in the tumor microenvironment is a relevant feature, as well, furthermore, it is important to analyze composition of the infiltrates [13]. The relation between effector T cells and regulatory T cells, among TILs, would have prognostic implications. To study regulatory T cells, Foxp3 should be considered as a reliable marker; consequently, the expression of this transcription factor was thoroughly analyzed in this study in an attempt to explore new potential immunotherapy targets. Therefore, the research of the immune modulation in BC, would not only help to prevent and reverse the resistance to conventional treatments but also, could highlight the development of new combined immunotherapies.

## MATERIALS AND METHODS

### Samples

Pre-treatment tumor samples from 122 breast cancer patients from different subtypes were collected from 2014 to 2019, during surgery, in hospitals related to the Faculty of Medical Sciences, National University of La Plata, Argentina. This research was approved by the Medical Bioethics Committee of the Institution (Protocol 41/2018). The World Medical Association Declaration of Helsinki (Finland, 1964) and further modifications were followed and Informed Consent was obtained from all patients included in this study. Clinical and histopathological data are summarized in Table 1. The mean age of the patients was 55 years old (range: 27-85 years old). Breast cancer samples were classified as: Invasive Ductal Carcinoma (IDC), Lobular Carcinoma (LC), Ducto-Lobular (DLC) and other types. Pathological staging was performed according to the 7th UICC-TNM.

**Table 1:**
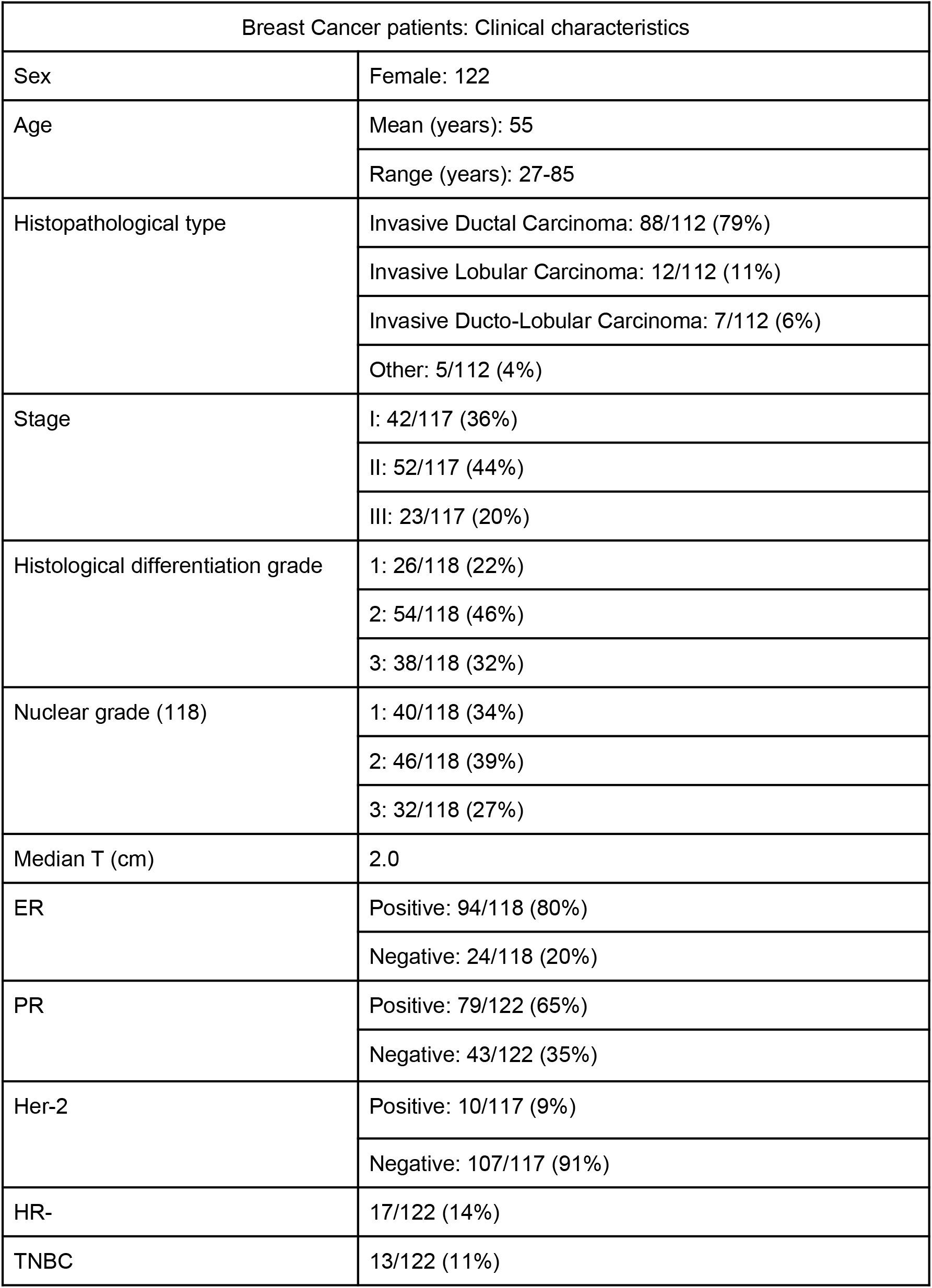
Clinical and histopathological characteristics of breast cancer patients

### Cell lines

Breast cancer cell lines MCF-7, ZR-75-1, T-47D and MDA-MB-231 were employed (ATCC).

### Primary antibodies

anti-IDO1 MAb (Clone 10.1, Chemicon-Millipore), anti-Foxp3 MAb (Clone ab22510, Abcam), anti-CD45RO MAb (BD) and anti-CD8 MAb (ThermoFisher) were employed at 1/50 dilution.

### Immunohistochemistry (IHC)

The technique was performed following standard procedures: FFPE specimens were treated with 10 mM sodium citrate buffer pH 6.0 at 100 °C for 10 min for antigen retrieval and incubated overnight at 4 °C with MAb. The reaction was developed with the LSAB+kit/HRP (DAKO, USA) following the manufacturer’s instructions. The chromogen employed was 3,3′-diaminodiazobenzidine (DAKO, USA). Sections were examined by light microscopy; the antibody staining patterns were scored in a semiquantitative manner. Staining intensity was graded as negative (0), low (1), moderate (2) or strong (3). The number of optical fields in a specimen that were positively stained was expressed as a percentage of the total number of optical fields containing tissue. The staining of the cytoplasm, plasma membrane and nucleus were evaluated; cells were considered positive when at least one of these components was stained. Sections were evaluated in a blinded way by two independent observers (MIL, MVC) following Croce et al. (2003) [14].

### Evaluation of TILs

Stromal and intratumoral infiltrating lymphocytes (sTILs and iTILs, respectively) were evaluated following Salgado et al. (2015) [13]. Briefly, the results were reported as the percentage of the area occupied by lymphocytes related either to total stroma (sTILs) or the tumor area (iTILs). TILs occupation percentages were classified in scores: negative (<10%); low (10-40%) and high (> 40%). Samples with more than 10% of the stromal or tumor area occupied by TILs (low or high scores) were TILs+.

### Cell culture

MCF7-, ZR-75-1, T-47D cell lines were grown in RPMI 1640 or DMEM, for MDA-MB-231 (SIGMA, USA), supplemented with 10% fetal bovine serum (Natocor, Argentina), 100 U/ml penicillin and 100 µg/ml streptomycin (P0781, Sigma-Aldrich).

### Immunocytochemistry (ICC)

Cell lines were grown over glass coverslips and fixed with 4% formaldehyde and a modified IHC was performed. The primary antibodies employed were anti-IDO and anti-Foxp3.

### In silico analysis

To evaluate the gene expression profile of *FOXP3* in the BC subtypes (Luminal A, Luminal B, ERBB2, Basal), the microarray dataset GSE21653 (n=266) and the RNAseq dataset TCGA PanCancer Atlas (n=1084) were employed. Triple Negative Breast Cancer (TNBC) subtypes were analyzed with the bioinformatics tool TNBCtype in GSE21653. Statistical analysis and plots were performed using the computing environment R and additional analysis packages from Bioconductor. To identify biological processes associated with *FOXP3* in BC, it employed *the guilt by association principle* based on gene expression profile similarities. Multiexperiment Matrix (https://biit.cs.ut.ee/mem/) was used to select the coexpressed genes (p<0.001; R>0.6) that correlate with *FOXP3* expression across 56 microarray datasets related to BC. Functional enrichment analysis was performed with the ClueGo plugin of the Cytoscape software to capture the biological processes associated with the *FOXP3* gene. Overrepresented GO terms, as well as <u>KEGG/Reactome/Wikipathways</u> pathways, were functionally organized into GO/pathway term networks.

### Statistical analysis

The analysis of Chi-square, univariate statistical analysis by Kendall’s correlation and multivariate analysis by PCA were performed employing IBM SPSS Statistics software. When p<0.05, statistical differences were considered significant.

## RESULTS

### Foxp3 and IDO1 expression in breast cancer samples are associated with tumor stage

Foxp3 and IDO1 expression was evaluated in BC samples. IDO1 expression was found in 76/122 (62%) of tumor samples while Foxp3 expression in 64/106 (60%). The immunohistochemical staining pattern was cytoplasmic and nuclear when samples reacted with anti-Foxp3 MAb (Figure 1A) and a mixed pattern (cytoplasmic and apical membrane) was found in samples showing reaction with anti-IDO1 MAb (Figure 1B).

**Figure 1.**
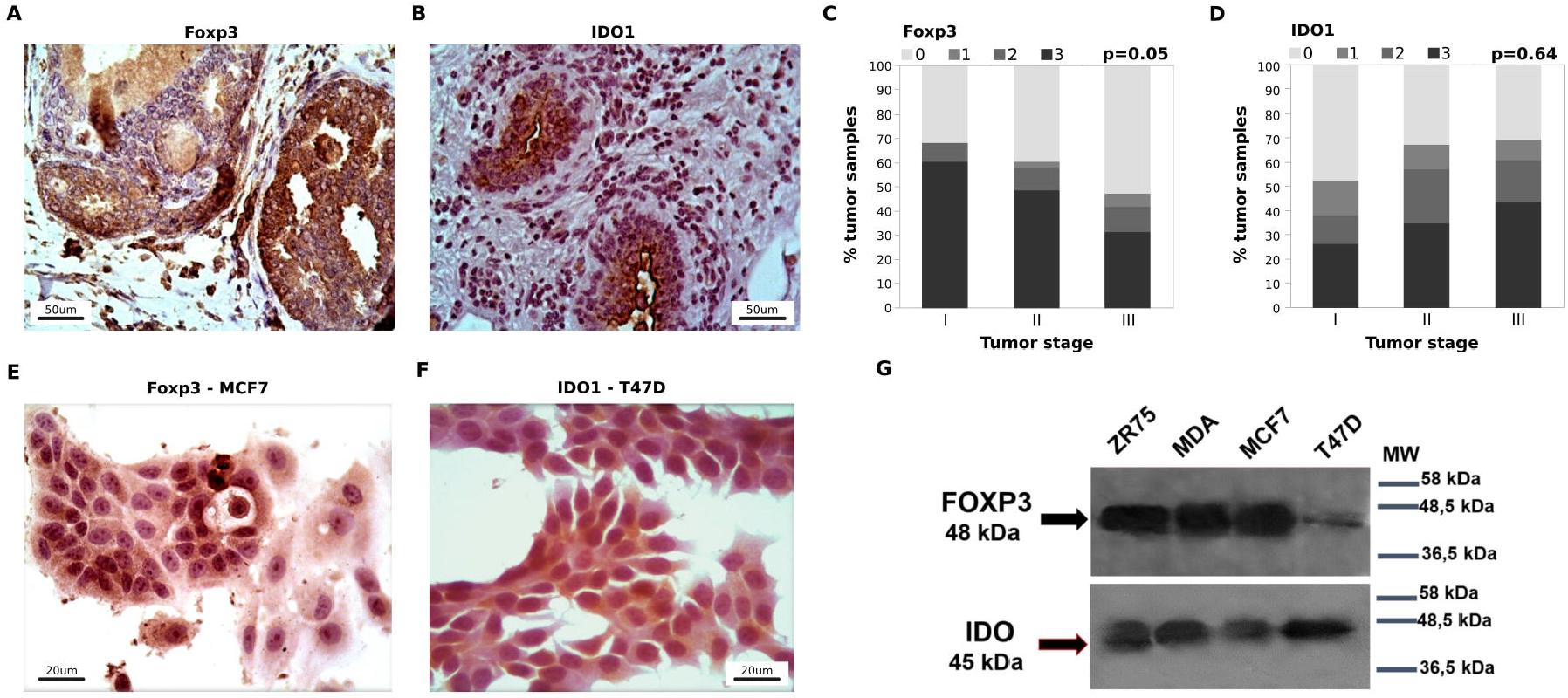
A. IHC of an IDC sample showing a strong Foxp3 expression with a cytoplasmic and nuclear pattern (100X). B. IHC of an IDC sample with a strong IDO1 protein expression with a mixed pattern (100X). C-D. Percentage of Foxp3+ or IDO1+ BC samples, with different intensities of expression according to TS. E-F. ICC of Foxp3 expression in MCF7 cell line and IDO1 in T-47D showing strong cytoplasmic and nuclear patterns of reaction (400X). G. Western blot showing Foxp3 (48 kDa) and IDO1 (45 kDa) expression in BC cell lines. Scale bar in micrometers.

When clinical and histopathological variables were included in the analysis, associations were found with tumor stage (TS), histologic grade (HG) and nuclear grade (NG).

A statistically significant negative correlation was detected between Foxp3 and TS. Increased Foxp3 expression was observed in early TS (stage I) in contrast to advanced TS (stage III) (p=0.05, Figure 1C). Conversely, high IDO1 expression was observed associated with advanced TS, although the association was not statistically significant (p=0.064, Figure 1D). Besides, a statistically significant positive association between IDO1 and NG was found (p=0.004), supporting previous results (Isla Larrain, 2014).

### Foxp3 and IDO1 expression in breast cancer cell lines

A panel of four BC cell lines: MCF7, ZR-75-1, T-47D and MDA-MB-231, was analyzed in order to detect Foxp3 and IDO1 expression. By ICC, Foxp3 showed a strong cytoplasmic and nuclear staining pattern, while IDO1 exhibited a cytoplasmic pattern (Figure 1 E-F). By western blot, Foxp3 (48kDa) and IDO1 (45kDa) protein expression were detected in the four cell lines studied (Figure 1 E-G).

### Foxp3 and IDO1 expression according to hormone receptor status

When samples were independently analyzed according to hormone receptor status, an interesting landscape of associations emerged.

ER negative tumors showed a statistically significant association between high Foxp3 expression and earlier TS (p=0.003, Figure 2A). In a similar way, lower HG was associated with high Foxp3 expression (p=0.001, not shown). In contrast, a statistically significant correlation was detected between high IDO1 expression and advanced TS (p=0.003, Figure 2B), as well as high HG (p=0.039, not shown).

**Figure 2.**
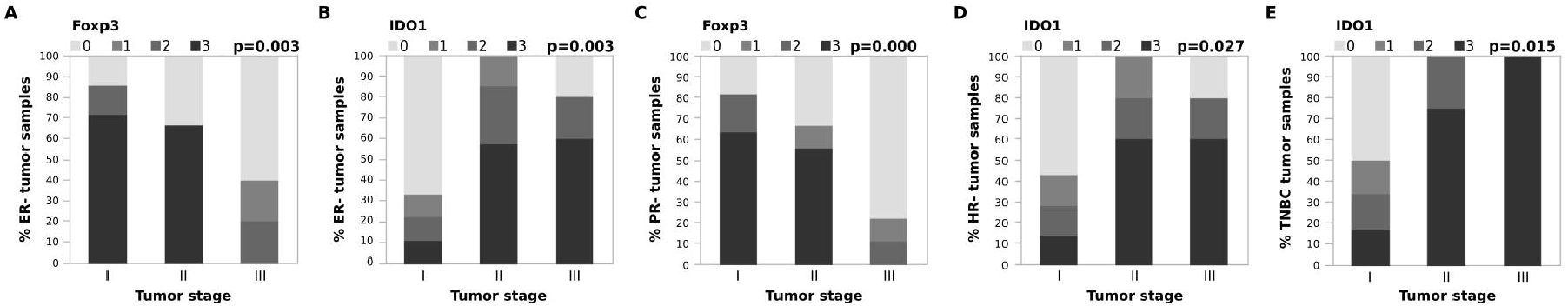
Foxp3 and IDO1 expression according to TS in ER-, PR-, HR- and TNBC tumor samples. A-B Percentage of Foxp3+ and IDO1+ samples in ER-tumors with different intensities of protein expression according to TS. C. Percentage of Foxp3+ samples in PR-tumors with different intensities of expression according to TS. D. Percentage of IDO1+ samples in HR-tumors with different intensities of protein expression according to TS. E. Percentage of IDO1+ samples in TNBC tumors with different intensities of expression according to TS.

PR negative tumors showed elevated Foxp3 expression at earlier TS (p=0.000, Figure 2C) and lower HG (p=0.000, not shown), while high IDO1 expression showed a non-statistically significant trend of association with advanced TS (p=0.066, not shown).

When tumor samples lacking ER and PR expression (HR-) were analyzed, a statistically significant negative correlation was observed between Foxp3 and IDO1 expression (p=0.014). Interestingly, all Foxp3 negative samples were IDO1 positive and showed higher HG (grade 3), besides 80% of samples corresponded to advanced TS. On the other hand, 55% of IDO1 positive samples showed none or low Foxp3 expression. Advanced TS tumors showed a statistically significant increment of IDO1 expression (p=0.027, Figure 2D), while a trend towards the loss of Foxp3 expression was observed (p=0.164).

Analysis of TNBC samples showed a positive statistically significant correlation between IDO1 expression and TS (p=0.015, Figure 2E). No associations were found in TNBC when Foxp3 expression and clinical or histopathological variables were analyzed.

### Tumor infiltrating lymphocytes landscape in breast cancer

The infiltration of lymphocytes in the microenvironment of tumor samples was also analyzed. Several characteristics of the TILs were evaluated: the distribution and the extent of the area occupied by the mononuclear cells, classified as intratumoral (iTILs) or stromal (sTILs). Scores were established following Salgado et al. (2015) [13]. Besides, the composition of TILs subsets was performed employing a-Foxp3 (a Treg marker), a-CD8 and a-CD45R0 (memory cell marker) MAbs.

Tumor infiltrating lymphocytes were found in 76% (93/122) of the tumor samples. Among the TILs positive samples, 42/93 (45%) showed exclusively iTILs; 7/93 (8%) exclusively sTILs; while 44/93 (47%) showed iTILs and sTILs concomitantly.

When sTILs were evaluated, a negative correlation with Foxp3 tumor expression was detected (p=0.000, Figure 3A). No association was found with IDO1 expression.

**Figure 3.**
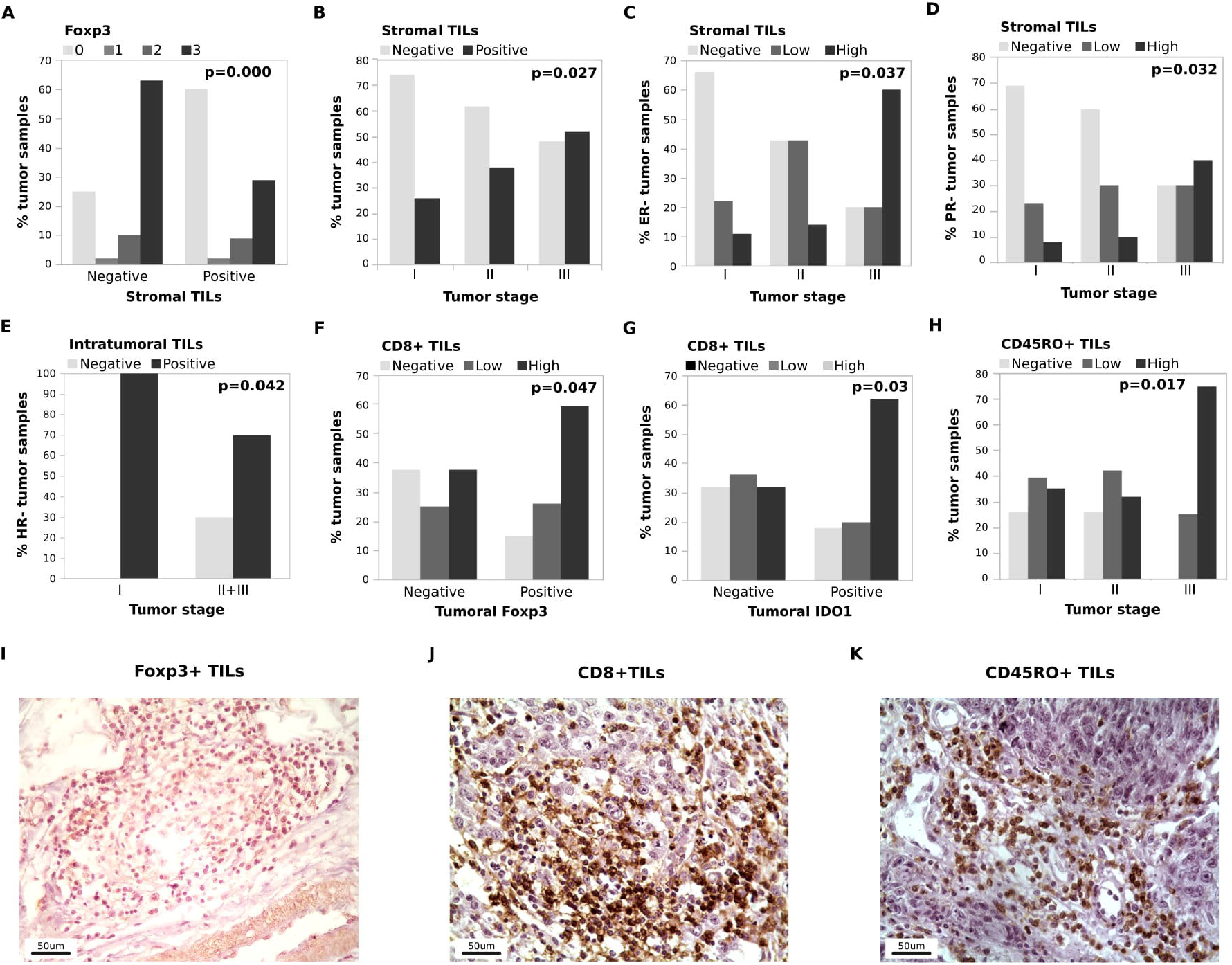
**A**. Percentage of BC samples expressing Foxp3 at different intensities vs. stromal TILs. B. Percentage of tumor samples with stromal TILs according to TS. C-D. Percentage of ER-samples or PR-samples with stromal TILs according to TS. E. Percentage of HR-samples with intratumoral TILs distributed by TS. F-G. Percentage of BC samples with CD8+ TILs in relation to Foxp3 or IDO1 tumor expression. H. Percentage of BC samples with CD45R0+ TILs according to TS. I-K. IHC of IDC samples showing Foxp3, CD8 and CD45R0 expression on lymphocyte infiltrates (400X). Scale bar in micrometers.

In relation to the clinical and histopathological variables, a positive association was detected between sTILs and TS. Stromal TILs were observed in 26% (11/42) of tumors classified as stage I, 41% (20/49) of stage II and 52% (12/23) of stage III (p=0.027, Figure 3B).

The association between advanced TS and increased sTILs in the tumor microenvironment was also observed when ER- and PR-samples were independently studied. Among ER-tumors, 60% of stage III samples showed high score sTILs in contrast to 11% of stage I samples (p=0.037, Figure 3C). Besides, when PR-tumors were analyzed, 40% of stage III samples showed high score sTILs in contrast to 8% of stage I samples (p=0.032, Figure 3D).

Concerning HG, a positive correlation with sTILs was found both in ER- and PR-groups (p=0.002 and 0.015, respectively; not shown). It means that poorly differentiated tumors showed increased lymphocyte infiltrate in the stroma in contrast to more differentiated tumors.

Regarding iTILs, a correlation with TNBC samples was found. In this BC subtype, 92% of samples showed iTILs in contrast with 69% of non-TNBC samples (p=0.014, not shown).

Among the HR-subset, a statistically significant negative correlation was found between TS and iTILs; 100% of stage I tumors exhibited iTILs, in contrast with 70% of advanced TS samples (p=0.042, Figure 3E).

The nature of lymphocyte infiltrate was also studied through the expression of Foxp3, CD8, and CD45R0.

Foxp3+ TILs were detected only in 15% of the TILs+ samples. On the other hand, 76% of samples showed CD8+ cells and 75% exhibited CD45R0+ cells.

Despite the limited number of samples, a statistically significant association was detected between Foxp3+ TILs and Foxp3+ tumors (p=0.000, not shown). Foxp3+ TILs were detected only in Foxp3+ tumors. Furthermore, most of these samples (92%) showed high tumoral Foxp3 expression, and were classified as TS I (67%), while the rest of samples were TS II and III, indicating an association between Foxp3+ TILs with early tumor stages.

On the other hand, abundant CD8+ TILs were found in tumor samples. Increased CD8+ TILs were observed in tumoral Foxp3+ samples (p=0.047, Figure 3F) and also in tumoral IDO1+ samples (p=0.030, Figure 3G). No statistically significant association was found when TS was considered, but higher HG and NG tumor samples showed increased CD8+ TILs (p=0.050 and 0.011, respectively; not shown).

When tumor samples were analyzed taking into account each hormone receptor, ER- and PR-tumors exhibited increased CD8+ infiltrate in contrast with ER+ and PR+ tumors (p=0.05 and 0.028, respectively; not shown). In contrast, no statistically significant associations for CD8+ TILs distribution were found between HR- and HR+ and between TNBC vs non-TNBC samples. However, 90% of HR- and 88% of TNBC tumors showed CD8+ TILs.

Regarding CD45R0+ cells, statistically significant correlations were found with advanced TS (p=0.017, Figure 3H), as well as with poorly differentiated BC samples (p=0.032, not shown). No other correlations were detected either with NG, tumoral Foxp3, IDO1 or hormonal receptor status. However, when HR- and TNBC samples were considered, 91% and 89% of samples, respectively, showed CD45R0+ cells infiltrate.

There was not found a statistically significant association between CD8+ and CD45R0+ tumor samples, however, 61% of the analyzed samples were positive for both markers.

Immunohistochemistry of Foxp3, CD8 and CD45R0 TILs positive samples are shown in Figure 3 I-K.

Suppl. file I shows the results of the different immunological variables analyzed according to receptor status.

### Basal and ERBB2/Her2-enriched tumor samples express higher levels of *FOXP3* than Luminal A samples

*In silico* analysis was performed to identify *FOXP3* coexpressed genes with Multiexperiment Matrix (p<0.001; R>0.6) across 56 microarray datasets related to BC (Suppl. file II). Functional enrichment analysis with the ClueGo plugin of the Cytoscape software was used to determine the biological processes associated with the *FOXP3* gene. Overrepresented GO terms, as well as <u>KEGG/Reactome/Wikipathways</u> pathways, were functionally organized into GO/pathway term networks (Suppl. file III).

*FOXP3* is upregulated in the ERBB2 and Basal intrinsic subtypes in contrast to Luminal A (p<0.001, Figure 4A). When histopathological variables were evaluated, it was observed that *FOXP3* is also upregulated in ER-, PR- and poorly differentiated tumors, in agreement with the features of Basal tumors (Figure 4B).

**Figure 4.**
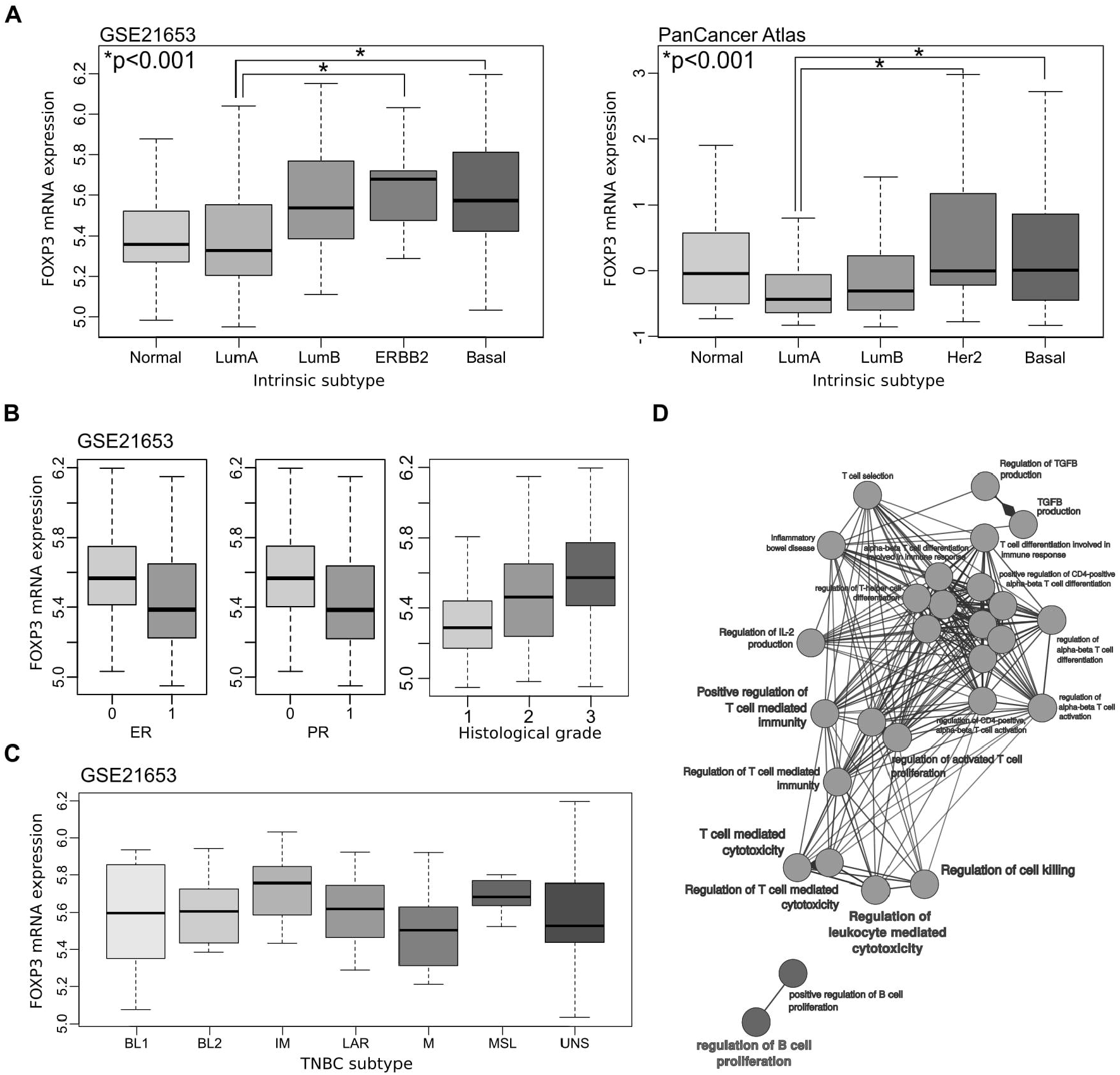
A. *In silico* analysis of *FOXP3* expression in BC subtypes employing the microarray dataset GSE2163 (n=266) and the RNAseq dataset TCGA PanCancer Atlas (n=1084). Basal and ERBB2 subtypes show significant differences with Luminal A in both datasets (p<0.001). B. *FOXP3* overexpression in ER-, PR- and undifferentiated tumors. C. *FOXP3* expression in the TNBC molecular subtypes. The highest expression level was found in the IM subgroup. No significant differences among groups were observed, although a tendency between M and IM was found (p=0.06). D. Functional enrichment analysis was performed with the ClueGo plugin of the Cytoscape software in order to capture the biological processes associated with the *FOXP3* gene co-expression network.

The bioinformatics tool TNBCtype was employed to classify *FOXP3* expression of Basal samples in the GSE21653 dataset according to the TNBC molecular subtype profiles proposed by Lehmann et al. (2011) [15]. TNBC samples are classified as Basal-like 1 and 2 (BL1 and 2), immunomodulatory (IM), mesenchymal (M), mesenchymal stem-like (MSL), luminal androgen receptor (LAR), and unstable (UNS). The highest *FOXP3* expression level was found in the IM subgroup, although this association was not statistically significant (Figure 4C). Functional analysis of *FOXP3* co-expressed genes was performed in BC datasets. Interestingly, the most relevant functional network cluster identified is associated with the biological term *regulation of leukocyte mediated cytotoxicity* and the ontological child categories: *regulation of activated T cell immunity, regulation of TGFB production*, among others (Figure 4D).

## DISCUSSION

The presence of immunomodulatory molecules and tumor infiltrating lymphocytes in BC microenvironment is critical in tumor dissemination and patient’s prognosis. In this study, the relationship between tumor cells and the microenvironment was performed by evaluation of several immune biomarkers such as IDO1, Foxp3, intratumoral and stromal TILs, CD8 and CD45R0 in breast cancer samples.

Both IDO1 and Foxp3 expression have been previously reported in tumors. Indoleamine-2,3-dioxygenase is an immunosuppressive enzyme with a protumorigenic effect [4]. IDO1 catalyzes the limiting step of tryptophan degradation through the kynurenine pathway. The decrease of the local concentration of tryptophan and the increase of the concentration of kynurenine in the IDO-expressing cells and nearby surrounding cells of the microenvironment exerts immunosuppressive functions through the regulation of T effector cells by anergy or apoptosis and the induction or activation of regulatory T cells [4, 16].

On the other hand, the role of Foxp3 in tumors remains controversial. This transcription factor is characteristic of regulatory T cells that exert immunosuppressive effects; however, it can be expressed by non-hematopoietic cells such as normal breast epithelium and has been reported as a suppressor gene or an oncogene in cancer biology [17, 18]. Foxp3 expression by tumor cells, as well as infiltration of Treg in tumor microenvironment, have been associated with different prognosis. In some neoplasms, such as colorectal cancer, melanoma, non-small and small lung carcinoma, high levels of Foxp3 were associated with poor prognosis. On the contrary, in breast, prostate and gastric cancer, high levels of Foxp3 were associated with good prognosis. This variability among different tumor locations is probably related to different functions of Foxp3 in given cases [19].

In a previous report, the role of IDO1 was studied in breast cancer. Overexpression of this enzyme was found in the cytoplasm of tumor cells and was associated with advanced tumor stages and higher nuclear grades but, in disagreement with Jacquemier et al. (2011), it was hardly found in normal mammary gland samples [5, 20]. In the current study, these results were confirmed, including the observation of IDO1 overexpression at ER-, PR- and TNBC tumors. Moreover, a positive correlation between IDO1 expression and histologic grade was observed. Interestingly, a statistically significant increase of IDO1 expression at advanced tumor stages in HR- and TNBC groups was found. In this sense, elevated *IDO1* gene and protein expression have been reported in TNBC tumors, as well as high IDO1 activity, in the serum of TNBC patients [21, 22].

Tumoral Foxp3 expression was also evaluated in BC samples; high levels of this transcription factor were found at early TS and a progressive loss of its expression was observed at advanced TS.

The associations of IDO1 and Foxp3 with opposite tumor stages were more evident when samples were independently evaluated according to the hormone receptors status. It is important to note that the inverse IDO1 and Foxp3 tumor expression according to TS was statistically significant in ER- and HR-tumors. Since both, IDO1 and Foxp3 are immunomodulatory variables, it could be suggested that their expression at different TS could be a compensatory mechanism of the tumor to maintain an immunosuppressive microenvironment.

Tumor infiltrating immune cells play a key role in tumor progression. Its ability to control tumor development and dissemination has been proved but is not fully understood. The study of TILs subsets and its distribution in relation to tumor cells in breast carcinoma is currently limited to the research field but is a promising biomarker for the development of more effective immune targeted therapy, playing a fundamental role in predicting the pathological complete response to neoadjuvant chemotherapy [23]. Since several studies have emerged with contradictory results about the effects of TILs in tumor progression and patients’ prognosis, choosing an appropriate therapeutic approach becomes difficult. [2, 3, 13, 23].

In this study, the distribution of TILs was evaluated in the tumor microenvironment for each sample, being classified as intratumoral or stromal. Both iTILs and/or sTILs were found in 72% of samples, indicating an ongoing antitumoral immune response. In general, a positive association between sTILs and advanced tumor stages was found. Regarding iTILs, a negative association with TS was observed in HR-tumors and TNBC samples showed only iTILs (not sTILs). Interestingly, all TNBC samples corresponded to advanced tumor stages. Several studies have reported that high levels of sTILs are associated with improved outcome and better response to neoadjuvant therapy in both TNBC and HER2-positive breast cancer [24]. Furthermore, in TNBC, sTILs can be used to identify a subset of patients at early TS with excellent prognosis without adjuvant chemotherapy [25]. Moreover, in a recent study, Vbihervuori et al. (2019) found that for every 10% decrease in sTILs there is a 20% increase in mortality risk for TNBC patients [26]. Therefore, the absence or low levels of sTILs in TNBC patients could be considered as a negative prognosis factor.

The evaluation of immune cell markers in tumor infiltrating lymphocytes showed a high frequency of CD8+ but not Foxp3+ TILs. This is very interesting given that Foxp3+ Treg are usually found in tumor infiltrates and in peripheral blood samples of BC patients, as well as other tumor locations [17]. Surprisingly, in this study Treg were hardly found in BC samples and when detected they were associated with Foxp3+ tumors in every case.

On the other hand, it was observed a high presence of CD8+ cells among TILs in tumor samples. The enrichment of CD8+ cells in the tumor microenvironment is an expected immune response. Cytotoxic CD8+ lymphocytes are hypothesized to be the final effector cells that mediate tumor cell killing, a mechanism of tumor destruction through recognition of tumor antigens and direct elimination of transformed cells [27, 28]. CD8+ are found in many solid tumors, therefore the presence and dynamics of CD8+ cells could be a reliable clinical marker and provide an attractive issue to be considered in immunotherapy [29]. It has been demonstrated a positive prognostic value of cytotoxic CD8+ TILs in breast cancer [30]. It is more, high CD4+TILs infiltration was revealed as an independent good prognostic factor in hormone receptor-negative subgroup while high Foxp3+/CD8+ cell ratio was found to be an independent adverse prognostic factor in hormone receptor-positive subgroup, especially in the luminal A subtype [31].

The positive correlation found in this study between tumoral IDO1 and Foxp3 expression with CD8+ TILs is very interesting. It could be suggested that while CD8+ TILs can be considered a good prognosis factor because its expected antitumoral activity, the expression of IDO1 and Foxp3 by tumors can be considered as bad prognosis factors since they can act as opposite forces that generate an immunosuppressive environment which counteracts the immune system’s attempt to eradicate the tumor.

The immunosuppressive effect of IDO1 was proved by Uyttenhove et al. (2003) [32] employing an IDO1-expressing murine model to show that a tumor with high IDO1 activity is not detected nor destroyed by tumor-specific host immune cells, caused by tryptophan (Trp) depletion. Similarly, Zaher et al. (2011) [33] demonstrated that downstream kynurenine pathway metabolites such as 3-hydroxykynurenine (Kyn), 3-hydroxyanthranilic acid (3HAA) and quinolinic acid can significantly inhibit T cell proliferation and induce T cell apoptosis. In this sense, several studies have described the immunomodulatory effects of IDO1 on CD8+ antitumor immune responses [34-36]. Moreover, CD8+ subsets described as exhausted, anergic, senescent and regulatory have been observed in clinical and basic studies [28]. On the contrary, [37], postulated that the constitutive upregulation of IDO1 expression may induce IDO1 specific T-cell responses that can directly recognize and kill IDO1+ tumor cells. Furthermore, it may even be speculated that the induction of IDO1-specific immune responses by therapeutic measures could function synergistically with additional immunotherapy not only by eliminating cancer cells but also immunosuppressive cells.

In relation to Foxp3 expression, its biological role in tumors is controversial. In TILs, this transcription factor has a crucial role in the generation of immunosuppressive CD4+ Treg cells and the induction of immune tolerance to antigens (<u>2</u>). However, it seems that Foxp3 in tumor cells may have distinct biological activities and prognostic values according to its cellular localization. Takenaka et al. (2013) suggested that Foxp3 cytoplasmic localization could hamper its biological role as onco-suppressor. Nuclear Foxp3 expression in tumor cells has been associated with improved overall survival in BC patients, whereas cytoplasmic Foxp3 has been associated with poor clinical outcome [17]. The nuclear role in transcriptional control of breast tumor suppressors (BRCA1) and oncogenes (HER2, SPK2) has been thoroughly analyzed [38, 38, 40]. In contrast, cytoplasmic Foxp3 and its functional role is not well understood. It has been proposed that Foxp3 cytoplasmic detection in cancer cells may result from defects in the nuclear localization signals due to acquired mutations, deletions, post-translational modifications and/or splicing variants [41, 42]. In this study, Foxp3 expression in breast carcinoma has been observed with a predominantly cytoplasmic pattern, and nuclear staining to a lesser extent. The main cytoplasmic localization of Foxp3 suggests that it could result from overexpression or could be caused by defects in the nuclear localization signals as it was mentioned above. By other side, it is important to point out that Foxp3 expression in tumor cells has been associated to inhibition of T cell proliferation and effector cytotoxic T lymphocyte (CTL) response, suggesting that Foxp3+ cancer cells may acquire immunosuppressive functions similar to Treg [43, 44]. Devaud et al. (2014) suggested a potential dual role of this transcription factor: on the one hand, it could be that a Treg-like activity by tumor cells is able to suppress effector T cells favoring tumor progression while, on the other hand, this transcription factor could act as a regulator of the inflammatory response, limiting tumor growth [18]. Recently, <u>Moreno-Ayala et al.</u> (2017) suggested a protumorigenic role for Foxp3. In another study, it has been reported that reprogramming of regulatory T cells to T helpers could enhance an effective CD8 response but remains unknown the spectrum of conditions under which Treg cells undergo reprogramming [45, 46].

It is important to note that in this study, a basal expression of Foxp3 was found in MCF-7, T47-D, MDA-MB-231 and ZR-75 cell lines. Has been described that the upregulation of Foxp3, following γ-irradiation in different human cell lines, was able to suppress the expression of the DNA-repair tumor suppressor gene BRCA1, a driver gene in breast cancer [40].

The expression of CD45R0, known as a marker of memory cells in adaptive immunity, was also evaluated in TILs. CD45R0+ cells display a low-activation threshold, which vigorously proliferates despite minimal co-stimulation and persists over a lifetime with stem cell-like multipotency and self-renewal characteristics [47]. In 75% of the BC samples included in this study, this memory T cell marker was detected at a high score (more than 40% of the tumor and/or stromal area occupied by these cells) mainly associated with advanced TS. It has been suggested that the presence of CD45R0+ effector cells in TILs is associated with a favorable prognosis for BC patients since they may both help eradicate local tumors and prevent lymphatic metastases. Besides, they also detected that a high ratio of CD45R0+/TILs was associated with recurrence-free survival improvement and a better cancer-specific survival [48].

In this study CD45R0+ cells were found in most of the tumor samples with CD8+ lymphocytes, suggesting that these TILs could be memory CTL (CD8+CD45R0+). In agreement with these results, Vahidi et al (2020) reported an increased frequency of CD8+CD45R0+ lymphocytes associated with large tumor size [49]. Furthermore, it has been reported that CD45R0 and CD8 expression and differentiation in immune cells parallelly increase in patients with BC malignant features such as advanced TS, poorly differentiated tumors and high number of nodes involved, as it was observed in this study. It would suggest that, despite the efforts of the immune system to provide an effective response to eradicate the tumor, it is often functionally impaired as a result of interaction with, or signals from, transformed cells and the tumor microenvironment [49]. These interactions and signals can lead to transcriptional, functional, and phenotypic changes that diminish their ability to eradicate the tumor [28].

The *in silico* analysis of *FOXP3* showed that Basal and ERBB2 tumor subtypes express higher levels of the transcription factor in relation to Luminal A subtype samples indicating that this transcription factor is associated with more aggressive subtypes. Interestingly, *IDO1* and co-expressed genes are also predominantly shown in Basal and ERBB2 subtypes as it was previously reported [5]. These findings suggest that both Foxp3 and IDO1 enhance immune escape, contributing to the tumor growth.

*FOXP3* expression was also evaluated in TNBC according to the immune molecular subtypes, with the result that the immunomodulatory subtype has the highest expression of the transcription factor, as it was also previously found for IDO1 expression [5]. It appears that IM characteristics are unique to the tumor cells themselves and not a reflection of TILs [15]. It has been established that the IM subtype is enriched for gene ontologies in immune cell processes, thus a functional analysis of *FOXP3* co-expressed genes was performed in BC datasets.

By *in silico* co-expression correlation analysis, *PD1CD1* standed up as one of the top *FOXP3* co-expressed genes. This gene codifies for the PD-1 immune checkpoint receptor, a well-known target of immunotherapy. This correlation would highlight the potential use of inhibition of Foxp3 in combined therapies. The immunotherapeutic potential of Foxp3 was suggested by Moreno Ayala et al. (2017) who employed the peptide P60 as an inhibitor of the transcription factor in BC cell lines [45]. The blockade of Foxp3 was associated with a decrease in IL-10 and TGFβ synthesized by tumors, implying that the expression of Foxp3 by the tumor could be involved in a mechanism of immune evasion. Besides, the most relevant functional network cluster identified from the list of co-expressed genes is associated with the biological term *regulation of leukocyte mediated cytotoxicity* and the ontological child categories: *regulation of T cell mediated immunity, regulation of TGF-B production*, among others. TGF-β signaling has been linked to the epithelial-mesenchymal transition and stem cell-like phenotypes exhibited by certain types of BC cells [50].

These findings are relevant in the context of the present study since it has been demonstrated that TILs have prognostic values in TNBC [51]. Thus, tumoral Foxp3 might constitute a relevant marker in TNBC tumors because of its association with TILs.

At the moment, the cumulative data suggest that Foxp3 could be a potential target in combined therapies, mainly in the most aggressive subtypes of breast cancer such as TNBC and Her2-enriched. In the same way, it has been suggested that using an IDO1 inhibitor in combination with a taxane (a chemotherapy agent which induces an influx of TILs in breast tumors) the ability of the effector cells to kill tumor cells may enhance clinical response. Indoximod (D-1MT), an IDO1 inhibitor, has shown to decrease IDO1-mediated suppression of the mTORC1 pathway in clinical trials [52]. Different approaches to direct IDO1 inhibition, including a peptide vaccine, have been promising and showed significant delay in tumor growth and prolonged survival in a B16 murine model. A variety of early clinical trials in different types of tumors are in progress, including IDO1 inhibitors with other checkpoint inhibitors such as anti-PD-1 and anti-CTLA4 [53]. Therefore, it should be relevant to perform the screening of IDO1 and Foxp3 in order to consider the inhibition of one or both biomarkers in combined immunotherapy.

Also, the nature and distribution of TILs should be considered when diagnosing and evaluating a patient’s treatment. To identify the composition and function of T cells in TILs, would have prognosis value for BC patients or improve the antitumor effect and therapeutic potential of adoptive cell therapy with TILs as was proposed by Sheng et al. (2017) for non-small cell lung carcinoma [24, 26, 54, 55].

## CONCLUSIONS

Foxp3 and IDO1 seem to have a relevant immunosuppressive role in the tumor microenvironment to counteract the host immune response characterized by high levels of CD8+ and CD45RO+ lymphocytes infiltrates. These two tumor markers could be emerging as the main immunosuppressive players at different tumor stages. Therefore, this research suggests the importance of performing a complete immunological profile of each BC patient through the evaluation of IDO1 and Foxp3 expression, as well as TILs distribution and composition, in order to design a suitable combined therapy to prevent resistance and relapse of the disease, mainly in aggressive subtypes such as TNBC and HER2-enriched.

## COMPLIANCE WITH ETHICAL STANDARDS

### Funding

This study was supported by the National University of La Plata (Research grant number M153-M185-M215) and the Comisión de Investigaciones Científicas de la Provincia de Buenos Aires (CICPBA)

### Ethical approval

All procedures performed in studies involving human participants were in accordance with the ethical standards of the Medical Bioethics Committee of the institution (2018) and World Medical Association Declaration of Helsinki (Finland, 1964) and its later amendments.

### Informed consent

Informed consent was obtained from all individual participants included in the study.

### Disclosure

Prof. María Virginia Croce, Dr. Marina Isla Larrain and Martín E. Rabassa are members of the Research Career of the Comisión de Investigaciones Científicas de la Provincia de Buenos Aires (CICPBA).

## Acknowledgements

We thank Mr. Federico Insaurralde for the technical assistance.

## LEGENDS TO THE FIGURES, TABLES AND SUPPLEMENTARY FILES

**Suppl. file I**. Results of IDO, Foxp3, sTILs, iTILs, CD8 and CD45RO expressed as positive samples/total (%) in the different subtypes of breast cancer samples according to hormone receptor status.

**Suppl. file II**. *In silico* analysis of *FOXP3* and coexpressed genes.

