## Supplementary file I for "Counterbalance of Foxp3 and IDO expression at different tumor stages in aggressive breast cancer subtypes"

**Suppl file I.** IDO, Foxp3, sTILs, iTILs, CD8 and CD45RO expressed as positive samples/total (%) in the different subtypes of breast cancer samples according to hormone receptor status.

| ER | PR | HER2 | Total | IDO | FOXP3 | iTILs | sTILs | CD8 | CD45RO |
|---|---|---|---|---|---|---|---|---|---|
| + | + | - | 66 | 37/66 (56%) | 35/57 (61%) | 43/66 (65%) | 20/66 (30%) | 22/32 (69%) | 31/43 (72%) |
| + | - | - | 19 | 12/19 (63%) | 9/17 (53%) | 12/19 (63%) | 7/19 (37%) | 9/9 (100%) | 13/13 (100%) |
| - | + | - | 5 | 4/5 (80%) | 4/5 (80%) | 4/5 (80%) | 2/5 (40%) | 4/4 (100%) | 2/4 (50%) |
| **Total ER+/- PR+/- HER2-** | | | **90** | **53/90 (59%)** | **48/79 (61%)** | **48/90 (53%)** | **29/90 (32%)** | **35/45 (78%)** | **46/60 (77%)** |
| + | + | + | 5 | 5/5 (100%) | 2/5 (40%) | 4/5 (80%) | 4/5 (80) | 2/5 (40%) | 5/5 (100%) |
| - | + | + | 2 | 1/2 (50%) | 1/1 (100%) | 1/2 (50%) | 0/2 (0%) | 1/1 (100%) | 0/1 (0%) |
| - | - | + | 4 | 2/4 (50%) | 2/3 (67%) | 3/4 (75%) | 2/4 (50%) | 1/1 (100%) | 2/3 (67%) |
| **Total ER+/- PR+/- HER2+** | | | **10** | **8/10 (80%)** | **5/9 (56%)** | **8/10 (80%)** | **6/10 (60%)** | **4/7 (57%)** | **7/8 (88%)** |
| **TNBC** | | | **13** | **10/13 (77%)** | **7/11 (64%)** | **12/13 (92%)** | **7/13 (54%)** | **7/8 (88%)** | **8/9 (89%)** |
